# How clonal are bacteria over time?

**DOI:** 10.1101/036780

**Authors:** B. Jesse Shapiro

**Affiliations:** Département de sciences biologiques, Université de Montréal, Montréal, QC H3C 3J7, Canada

## Abstract

Bacteria and archaea reproduce clonally (vertical descent), but exchange genes by recombination (horizontal transfer). Recombination allows adaptive mutations or genes to spread rapidly within (or even between) species, and reduces the burden of deleterious mutations. Clonality – defined here as the balance between vertical and horizontal inheritance – is therefore a key microbial trait, determining how quickly a population can adapt and the size of its gene pool. Here, I discuss whether clonality varies over time and if it can be considered a stable trait of a given population. I show that, in some cases, clonality is clearly not static. For example, non-clonal (highly recombining) populations can give rise to clonal expansions, often of pathogens. However, an analysis of time-course metagenomic data from a lake suggests that a bacterial population’s past clonality (as measured by its genetic diversity) is a good predictor of its future clonality. Clonality therefore appears to be relatively – but not completely – stable over evolutionary time.

## Introduction

Here, I revisit the question posed in the title of a classic paper by John Maynard Smith and colleagues [1]: How clonal are bacteria, and more specifically how does clonality vary among different microbial populations and over time? First, what do we mean by clonality? Perfectly clonal bacteria replicate by cell division (vertical descent) and evolve by random mutations that occur during DNA replication. In this theoretical population, there is negligible horizontal transfer of DNA by recombination across the resulting tree of vertical descent. Very few (if any) natural bacterial populations fit this idealized, theoretical definition of clonality. Or, as discussed below, they might only fit it for a short amount of time. However, knowing where a bacterial population of interest happens to fall along a spectrum of clonality can help us understand its biology, and even make predictions about its evolution.

The opposite of clonality is panmixis – a situation in which the rate of horizontal transfer is higher than the rate of vertical cell division, resulting in random association (linkage equilibrium) among loci in the genome [1,2]. However, rates of horizontal transfer (recombination) vary widely across the genome, such that a population can be mostly clonal, except for a few loci in the genome [3]. These loci came to be termed genomic islands – a metaphor I will build upon below. Some of the first islands identified were called pathogenicity islands because they contained virulence factors [4]. However, non-pathogenic environmental bacteria also contain islands, conferring adaptation to different ecological niches. For example, genes in *Prochlorococcus* genomic islands confer adaptation to light and nutrient conditions [5,6]. But islands need not confer niche adaptation to their host genome; they can be neutral to host fitness or even detrimental, selfish parasites. Here, I define genomic islands broadly as any piece of DNA that is transferred horizontally (by either homologous or nonhomologous recombination) from cell to cell and therefore evolves independently (*i.e*. is unlinked) from the rest of the genome.

I will begin by extending the use of island analogies to include continents, peninsulas and archipelagos (Table 1). I will then use these analogies to discuss to what extent microbial populations are clonal or panmictic, and how often they transition between the two regimes.

**Table 1.**
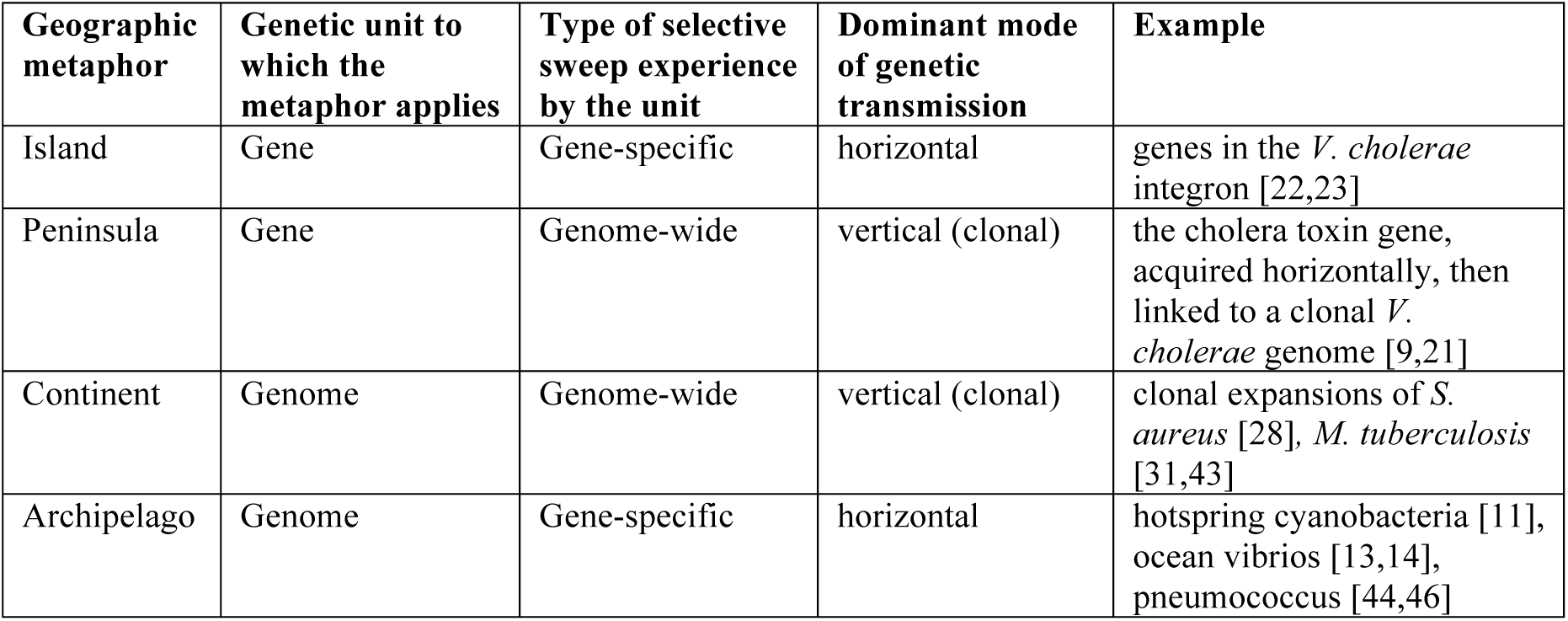
Extended island metaphors of microbial genome evolution.

| Geographic metaphor | Genetic unit to which the metaphor applies | Type of selective sweep experience by the unit | Dominant mode of genetic transmission | Example |
| --- | --- | --- | --- | --- |
| Island | Gene | Gene-specific | horizontal | genes in the *V. cholerae* integron [22,23] |
| Peninsula | Gene | Genome-wide | vertical (clonal) | the cholera toxin gene, acquired horizontally, then linked to a clonal *V. cholerae* genome [9,21] |
| Continent | Genome | Genome-wide | vertical (clonal) | clonal expansions of *S. aureus* [28], *M. tuberculosis* [31,43] |
| Archipelago | Genome | Gene-specific | horizontal | hotspring cyanobacteria [11], ocean vibrios [13,14], pneumococcus [44,46] |

## Are some islands really peninsulas?

In the classic analogy, an island is totally disconnected from the mainland, meaning that genes in the island evolve independently of the genome (Table 1). Examples of islands that fit this strict independence might include integrated phages and other “selfish” elements, or genes that reside in a particular niche but not in a particular genome (*e.g*. a gene ecology model [7]). Peninsulas provide an analogy that might better describe how islands are related to microbial genomes. A peninsula (or "presque-ile," from the French for "almost island") is a geographic term for a very narrow strip of land connected to (but distinct from) the mainland. In my analogy, an island is evolutionarily independent of the mainland genome, but their fates may become linked, forming a peninsula. For example, a bacterium may acquire a gene from a vast microbial gene pool. This gene allows the bacterium to invade a new ecological niche, triggering a clonal expansion in which the fate of the gene and its new host genome are linked, at least for the duration of the clonal expansion. One such example could be *Yersinia pestis*, which acquired a single gene allowing flea-borne transmission and triggering a clonal expansion in the form of Plague pandemics [8]. Another peninsula, the prophage-encoded cholera toxin, and its links to the mainland *Vibrio cholerae* genome [9,10], is discussed below.

## Are some genomes archipelagos?

The very concept of one or a few islands implies a contrast with the large, clonal genomic mainland or continent. But some microbial genomes may contain so many islands that there is no mainland, only a vast archipelago (Table 1). A striking recent example is a population of hotspring cyanobacteria in which virtually every gene in the genome evolved independently due to frequent recombination [11], leading the authors to call the population “quasi-sexual” (in other words, panmictic). Frequent recombination was confirmed by another group, using different methods to study the same cyanobacteria [12]. This group also found that despite a history of panmixis over long time scales, populations are more clonal over shorter time scales. Similarly, the Asian ocean population of *Vibrio parahaemolyticus* also forms a panmictic gene pool, with each recombination block of ~1.8 kbp evolving independently [13]. However, the panmictic gene pool occasionally gives rise to pandemic clones. In another example, we found that almost every gene in a population of *Vibrio cyclitrophicus* genomes showed signs of recombination over relatively recent time scales [14]. Such apparently high rates of recombination in natural populations were mysterious at first, contradicting recombination rates measured in the lab [15,16] and predicted by theory [7,17]. However, theoretical models (discussed below) suggest mechanisms capable of explaining how genes can spread through populations more rapidly by recombination than by clonal expansion [18–20].

## Clonal expansions from panmictic pools

I propose that archipelagos are not necessarily static over time, and that archipelagos can sometimes coalesce into continents. Given the right ecological opportunity, a genome from a panmictic gene pool can escape the "gravitational pull" of recombination and take off into a clonal expansion. An example mentioned earlier is *V. cholerae*, a genetically diverse group of coastal marine bacteria, some of which cause cholera. Virulence is mainly determined by two loci in the genome: the cholera toxin and the toxin-coregulated pilus. Both genes are frequently gained and lost by recombination [21,22], but are always found in one lineage of *V. cholerae* – the lineage causing severe disease with pandemic potential, known as the phylocore genome (PG) group [10]. It remains a mystery why the PG lineage evolved once, and only once. If PG *V. cholerae* really did evolve just once, this would be surprising because *V. cholerae* draws on a diverse, global gene pool and can be considered panmictic [23]. Therefore multiple different lineages would be expected to acquire the two (or perhaps a handful of) genetic elements required for pandemic disease. This leads to the hypothesis that pandemic cholera emergence is *selection limited* rather than *diversity limited*. In other words, benign *V. cholerae* strains constantly acquire virulence genes. However, these strains rarely encounter the right ecological niche to flourish, *e.g*. a human population consuming brackish water. "The right niche" has appeared a few times in human history: for example in India in the 1800s, when the Classical lineage evolved, and again in Indonesia in the 1950s, when the El Tor lineage evolved [24]. When the right conditions appear, the PG lineage, along with its virulence factors, takes off in a clonal expansions which continue to wreak havoc today (*e.g*. cholera pandemics from the 1800s to today, all caused by the PG clonal group). The virulence factors, previously islands in an archipelago, became a peninsula connected to the PG mainland. The linkage between virulence factors and PG remains imperfect because different variants of the cholera toxin continue to flow in and out of the PG continent [10,21]; hence the toxin remains a peninsula, not firmly part of the mainland.

*V. cholerae* is a particularly well-characterized example of a panmictic gene pool giving rise to a clonal expansion, but similar evolutionary dynamics are seen in other pathogens as well (*e.g. V. parahaemolyticus* described above [13]). Enterotoxigenic *Escherichia coli* (ETEC) seems to behave similarly, with deep branches of the phylogeny obscured by frequent recombination and plasmid exchange, but more recent branches experiencing mostly clonal descent, with tight linkage between virulence factors and the genomic mainland [25]. These observations are consistent with an ancient, panmictic gene pool giving rise to clonal expansions, which can last for decades or centuries.

## The balance between recombination and selection

Let us consider the evolutionary forces that determine clonality: natural selection and recombination. The effect of recombination on clonality is straightforward: more recombination means less clonality. The effect of natural selection is more complex, but is defined here simply as a force which favors clonal expansions of adaptive mutants within an ecological niche. Selection, as defined here, therefore includes ecological effects. When driven by ecological selection, clonal expansions are called selective sweeps, in which one clone outcompetes all others, purging genetic diversity in the population.

Recombination and selection interact to determine the clonality of a population. Recombination rates depend both on the ability of DNA to enter a cell and be incorporated into the genome (the baseline rate) and the ability of that DNA to be retained by a balance of genetic drift and natural selection (the realized rate). Some bacteria, such as *Helicobacter pylori,* have realized recombination rates that are much higher than point mutation rates, exchanging at least 10% of their genome within a single four-year human infection [26]. Others, such as *Staphylococcus aureus* [27,28] and *Mycobacterium tuberculosis* are decidedly more clonal [29–31]. Recombination rates (both realized and baseline) vary widely across the genome. Of 10 pathogenic bacterial species studied, all had identifiable recombination 'hot' regions, although their length, genomic location and gene content varied [3]. Genes of different functions had different realized recombination rates, implying a role for natural selection on gene function in determining whether newly acquired genes are retained.

## Modeling the recombination-selection balance

When rates of recombination are relatively low compared to rates of natural selection on adaptive genes within niches, entire genomes will sweep to fixation before they can be shuffled by recombination. Following previous modelling work, *s* is defined here as the selective coefficient of a niche-adaptive allele, and *r* is the recombination rate, per locus per generation [7,32,33].The *s* >> *r* regime is well described in the Stable Ecotype Model [17], which predicts that most of the genome will follow a single, clonal phylogeny. Genome-wide sweeps thus increase clonality and can be considered a hallmark of clonal populations (Table 1, Figure 1A). In the *r* >> *s* regime, individual genes (rather than entire genomes) will sweep to fixation (*i.e*. reach 100% frequency) in ecological niches to which they are adapted, without affecting genetic diversity elsewhere in the genome (Figure 1B). The first serious theoretical attempt to reconcile the observations of gene-specific sweeps with low recombination rates was the "Adapt Globally, Act Locally" model [18,20], in which globally adaptive genes (adaptive in multiple different niches) trigger genome-wide sweeps within a niche before being transferred to the next niche. When viewed across niches, the result is a gene-specific sweep. A recent model suggested another mechanism by which these gene sweeps can occur at moderate (not unrealistically high) rates of recombination [20]. In this model, a microbial habitat is bombarded with genetically maladapted migrants, allowing gene sweeps to occur, although the adaptive allele never reaches 100% frequency due to the constant input of migrants. In another model, Takeuchi et al. [19] show that gene sweeps can occur when *r* is either very high or – counter-intuitively – when *r* is very low, but only when negative frequency-dependent selection (NFDS) reduces the rate of genome-wide selective sweeps. NFDS might be commonly imposed on bacteria and archaea by viral (phage) predation, providing a selective advantage to rare alleles of phage receptor genes [34].

**Figure 1.**
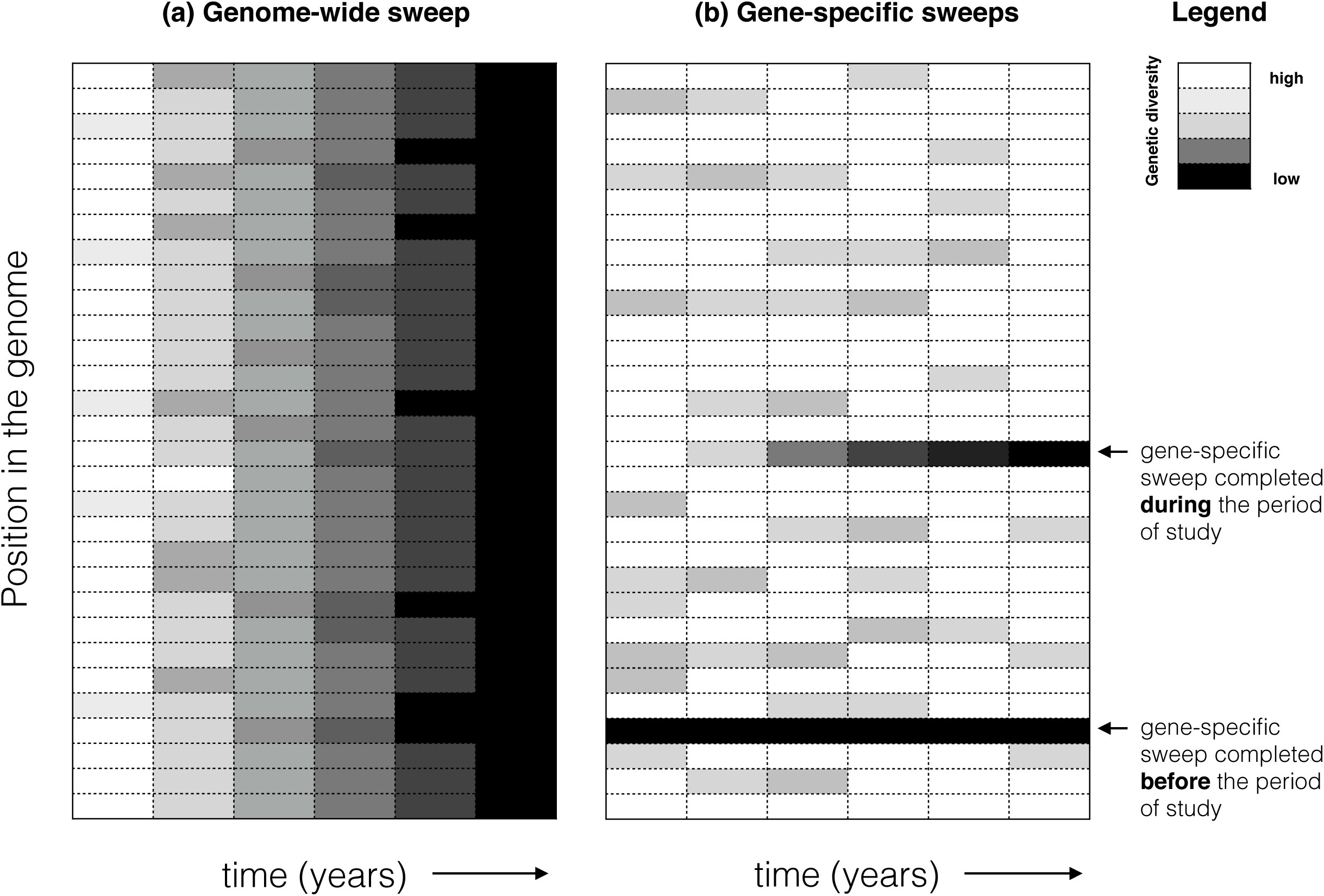
Temporal dynamics of genome-wide and gene-specific selective sweeps inferred from metagenomic data. Genetic diversity can be measured by mapping metagenomic reads to a reference genome, identifying SNPs, and calculating the allele frequencies at each SNP position in the genome over time. The lowest possible genetic diversity occurs when a single allele is present in 100% of metagenomic reads. Alternatively, diversity could be defined in terms of gene presence/absence, based on relative coverage of a gene in the reference genome by metagenomic reads. (**a**)In a hypothetical genome-wide selective sweep, all positions in the genome tend toward low diversity. (**b**)In a hypothetical gene-specific selective sweep, only one or a few positions in the genome tend toward low diversity, while the rest of the genome maintains high or intermediate diversity.

One parameter not thoroughly explored in any of these models is the effective population size (N_e_). Populations with small N_e_ are dominated by drift, and natural selection is inefficient. They may experience genome-wide sweeps independently of *r* and *s*. As discussed below, N_e_ is probably an important determinant of clonality.

## Genome-wide and gene-specific sweeps in nature

To date, empirical evidence for gene-specific and genome-wide sweeps has come mostly from cross-sectional studies of a single population of genomes at a single point in time, with recombination and selection inferred backward in time [11,13,14,35]. Sequencing microbial genomes or metagenomes sampled over time – already a typical practice in genomic epidemiology (e.g. [28,36]) – promises to elucidate the rates of gene-specific and genome-wide sweeps in nature ( Figure 1).

In a pioneering study, Bendall et al. [37] sampled a lake over nine years and followed single-nucleotide polymorphism (SNP) and gene frequencies in 30 bacterial populations by metagenomic sequencing. They inferred that one of the populations (*Chlorobium*-111) had undergone a near-complete genome-wide sweep over the nine-year study, with most SNP diversity purged genome-wide (Figure 1A). By identifying regions of the genome with unexpectedly low diversity compared to the genome-wide average, they inferred that genes-pecific sweeps had occurred in six of the populations, but these sweeps occurred before the beginning of the nine-year study. During the nine-year study, they observed examples “where a few adjacent SNPs trended toward fixation while genome-wide diversity was maintained” (Figure 1B). They took this observation as consistent with gene-specific selective sweeps, but did not attempt to determine whether the sweeps were due to selection or drift. Similarly, the inferred genome-wide sweep could have been caused by selection or drift.

Is the frequency of genome-wide sweeps controlled mainly by the balance of recombination and selection, or could drift (controlled by the effective population size, N_e_) play an important role as well? In their Figure 2 legend, Bendall et al. observe that "populations with many SNPs were not necessarily sequenced deeper than those with few SNPs." This statement was simply meant to assure the reader that SNP calling was not biased by sequence coverage, but it also suggests that microbial populations tend to be far from a standard neutral model with constant population size (*i.e*. the Wright-Fisher coalescent [38]). Taking SNP density as a rough measure of Ne and sequence coverage as a rough measure of the census population size, the data show that census population size is a poor predictor of N_e_, and thus that populations are likely impacted by population bottlenecks and/or selection – although it is difficult to distinguish between them.

**Figure 2.**
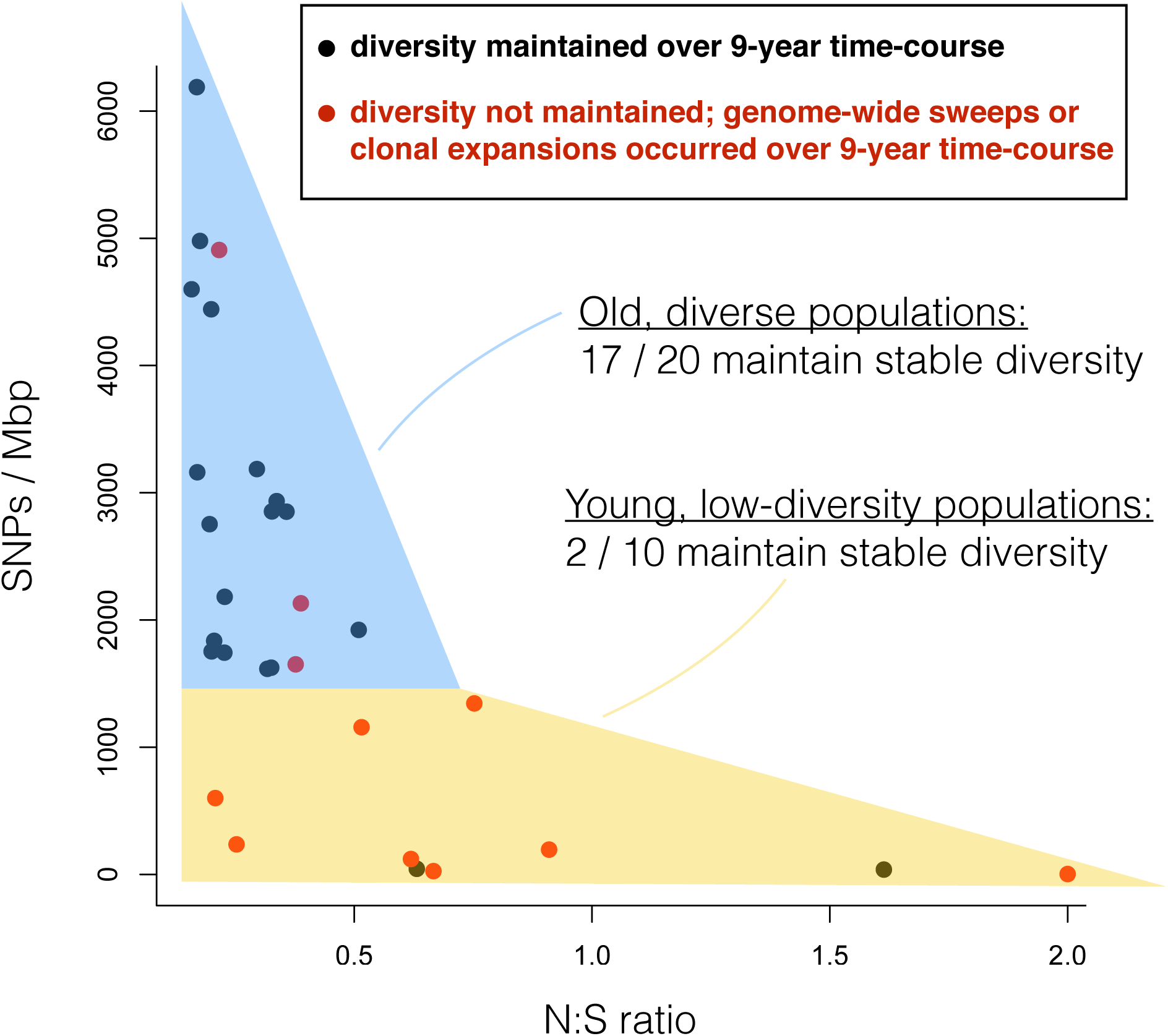
Past diversity predicts future diversity. Based on data from Table 2 of Bendall et al. [37], the nonsynonymous to synonymous (N:S) SNP ratio was plotted against the total SNP density (SNPs per megabasepair) for each of 30 bacterial populations. A pseudocount of 1 was added to both N and S counts. These 30 populations were divided into 20 “old, diverse” populations (>1500 SNPs/Mbp) and 10 “young, low-diversity” populations (<1500 SNPs/Mbp). Each point represents one of the 30 populations, colored in black if diversity was maintained over a 9-year metagenomic time-course, or in red if it was not. Seventeen out of 20 “old, diverse” populations maintained their diversity over the 9-year study, compared to only 2 out of 10 “young, low-diversity” populations (Fisher test, Odds Ratio = 19.4, *P* < 0.001). The same result is obtained drawing a cutoff based on N:S rather than SNPs/Mbp: populations with N:S < 0.5 tend to maintain their diversity, whereas those with N:S > 0.5 tend to be purged of diversity over the 9 years (Fisher test, Odds Ratio = 5.94, *P* = 0.042). Consistent with previous observations that N:S depends on the evolutionary time available for purifying selection to act [47,48], N:S is negatively correlated with SNPs/Mbp, a proxy for evolutionary time or the time since the last genome-wide purge of genetic diversity in this dataset (Pearson’s correlation of log_10_ transformed data, *r* = −0.81, *P* = 5.6e-8).

As a whole, the study showed that both genome-wide and gene-specific sweeps can occur in different microbial populations from the same environment. Whether microbial populations behaved differently due to differences in their ecology (*i.e*. regime of natural selection) or in their baseline recombination rates remains a question for future study; Cohan recently suggested that ecological differences could be more important [39]. Specifically, he suggests that ecological "generalists" could have more opportunities for diversification, and thus be relatively immune to genome-wide sweeps, compared to ecological specialists. I suggest that these generalists may simply have larger N_e_, such that diversity is rarely purged by drift (*e.g*. bottlenecks), and that diversity is mainly governed by selection and recombination. It is possible that ecological generalists tend to have large N_e_, or that N_e_ and ecology exert independent effects (*e.g*. ecological generalists can have low N_e_ and still resist genome-wide sweeps). The fact that one genome-wide sweep was observed over a nine year period suggests that such events might be relatively rapid but rare (only observed in one of 30 populations). Meanwhile, gene-sweeps might be more common historically (affecting six of 30 populations), but could take longer to proceed to completion.

## Is clonality a stable trait?

As described in the *V. cholerae* example, some pathogenic bacterial populations can switch between panmictic and clonal lifestyles [11,13,14,19,25,34,35]. Therefore clonality can vary over time, but how much and how often? To quantify the stability of clonality over time, not just in pathogens but in free-living environmental bacteria, I re-analyzed the lake time-course of Bendall et al. [37]. Because estimates of selection and recombination rates were not readily available for this dataset, I defined clonality based on the total genetic diversity (SNP density) in a population, which scales with N_e_, and includes the effects of both drift and of genome-wide selective sweeps. Frequent genome-wide selective sweeps (*s* >> *r*) and/or small N_e_ result in clonality. I identified 20 “old, diverse” populations as those with a high density of SNPs (>1500 SNPs/Mbp) at the beginning of the time-course. These populations are likely “old” because they have gone a relatively long time since the last genome-wide purge of genetic diversity and are relatively non-clonal. They include the six populations inferred to have undergone gene-specific sweeps [37]. The remaining ten populations were defined as “young, low-diversity,” having more recently experienced a genome-wide purge of diversity. The “old, diverse” populations have a relatively low ratio of nonsynonymous (N) to synonymous (S) SNPs, suggesting large N_e_ and ample time for purifying selection to remove (mostly deleterious) nonsynonymous mutations (Figure 2). In contrast, the “young, low-diversity” populations are more likely to have high N:S ratios, suggesting smaller N_e_ and less time for purifying selection to act. The cutoff of 1500 SNPs/Mbp (0.15% SNP density) is somewhat arbitrary, but seems to correspond to an inflection point in Figure 2, and also falls squarely within the 0-0.30% SNP density, above which *E. coli* evolution appears to transition from clonal to recombining [40].

With young (presumably clonal, or low N_e_) and old (less clonal, higher N_e_) populations thus defined, I asked whether “old, diverse” populations tended to maintain their diversity through the 9-year period of the study. In their paper, Bendall et al. defined two alternative population types: 1) those that maintained stable SNP diversity over nine years, and 2) those that experienced significant fluctuations in diversity due to clonal expansions – defined when one, but not all timepoints are dominated by a single allele (≥95% frequency) at >40% of SNP sites in the genome [37]. By definition, the 19 populations of the first type did not experience genome-wide sweeps during the study, while the 11 populations of the second type did experience genome-wide purges of diversity, which were transient in ten cases and apparently permanent in one case (*Chlorobium*-111) Strikingly, 17 out of 20 “old, diverse” populations maintained their diversity over the nine-year study, compared to only two out of ten “young, low-diversity” populations (Fisher test, Odds Ratio = 19.4, *P* < 0.001). This result suggests that populations with a history of genome-wide sweeps tend to experience subsequent genome-wide sweeps, and those that have maintained genetic diversity in the past tend to maintain their diversity into the future. In other words, clonality can be considered a relatively stable microbial trait. However, we must take care in taking a genome-wide sweep as evidence for clonality (*s* >> *r*). In small effective population sizes, genome-wide sweeps can occur due to drift and bottlenecks, independently of *s* and *r*. Therefore, many of "young, low diversity" populations (Figure 2) may have lost diversity over nine years due to low N_e_, not due to low *r*.

## History repeats itself

It appears that pathogens are more likely than free-living bacteria to undergo clonal expansions, due in part to their ecology and transmission dynamics [41]. Free-living aquatic bacteria, on the other hand, seem to be more likely to live in large, panmictic populations and behave like archipelagos [11,13,14,37]. If clonality is indeed a stable trait, this implies that history will repeat itself, and that the future behavior of microbial populations can be predicted with some confidence from their past behavior. Diverse populations tend to stay diverse. Clonal populations (that experience frequent genome-wide sweeps) tend to stay clonal. But history is not doomed to repeat itself forever. As we have seen, clonal expansions, such as pandemic *V. cholerae*, may originate when a panmictic gene pool (an archipelago) coalesces into a clonal continent, with virulence factors linked as peninsulas. Many such pathogenic clones have been documented, with life spans of decades to thousands of years [25,28,42,43]. Other pathogens, such as *Streptococcus pneumoniae*, may retain their panmictic population structure throughout an outbreak [44–46]. Why some pathogens are clonal and others are panmictic is an open question, but surely depends on the balance between recombination and selection, on the effective population size, and on the time scales considered.

## Acknowledgments.

I am grateful to Rex Malmstrom, Yan Boucher, and Salvador Almagro-Moreno for their thoughtful comments which improved the manuscript. I was supported by a Canada Research Chair.

## Footnotes

* A comprehensive discussion of the roles of selection and recombination in structuring microbial diversity.

* Deep sequencing of 90 cyanobacterial marker genes reveals extensive recombination, with each genome composed of a random mix of alleles from the gene pool.

* In-depth population genomic evidence of a panmictic ocean gene pool containing non-clonal ecological populations.

* Mathematical modeling shows how individual genes can sweep through populations with low recombination rates in the presence of negative frequency-dependent selection.

* A clear demonstration that genes in the integron are recombined between species just as often as within species, and that 24% of ‘core’ genes have crossed species boundaries.

** A pioneering application of time-course metagenomics to infer genome-wide and gene-specific selective sweeps in natural lake bacterial populations.

* The first simulation of bacterial evolution including mutation, allelic exchange (homologous recombination) and gene gain/loss (non-homologous recombination).

